# CLPX regulates erythroid heme synthesis by control of mitochondrial heme synthesis enzymes and iron utilization

**DOI:** 10.1101/2020.09.09.289660

**Authors:** Catherine M. Rondelli, Mark Perfetto, Aidan Danoff, Hector Bergonia, Samantha Gillis, Gael Nicolas, Herve Puy, Richard West, John D. Phillips, Yvette Y. Yien

## Abstract

Heme is a prosthetic group that plays a critical role in catalyzing life-essential redox reactions in all cells, including critical metabolic processes. Heme synthesis must be tightly co-regulated with cellular requirements in order to maximize utilization and minimize toxicity. Terminally differentiating erythroid cells have an extremely high demand for heme for hemoglobin synthesis. While the enzymatic reactions of heme synthesis are extremely well studied, the mechanisms by which the mitochondrial homeostatic machinery interacts with and regulates heme synthesis are poorly understood. Knowledge of these regulatory mechanisms are key to understanding how red cells couple heme production with heme demand. Heme synthesis is tightly regulated by the mitochondrial AAA+ unfoldase CLPX, which has been reported to promote heme synthesis by activation of yeast δ-aminolevulinate synthase (ALAS/Hem1). CLPX was also reported to mediate heme-induced turnover of ALAS1 in human cells. However, a mutation in the ATP binding domain of CLPX that abrogated ATP binding caused an increase in ALAS activity, contrary to previous predictions that CLPX activated ALAS. Using loss-of-function assays in murine cells and zebrafish, we interrogated the mechanisms by which CLPX regulates erythroid heme synthesis. We found that consistent with previous studies, CLPX is required for erythroid heme synthesis. We show that ALAS2 stability and activity were both increased in the absence of CLPX, suggesting that CLPX primarily regulates ALAS2 by control of its turnover. However, we also showed that CLPX is required for PPOX activity and maintenance of FECH levels, likely accounting for the heme deficiency in the absence of CLPX. Lastly, CLPX is required for iron metabolism during erythroid terminal differentiation. Our results show that the role of CLPX in heme synthesis is not conserved across eukaryotes. Our studies reveal a potential mechanism for the role of CLPX in anemia and porphyria, and reveal multiple nodes at which heme synthesis is regulated by the mitochondrial housekeeping machinery.

## Introduction

Heme is a prosthetic group that consists of a central iron chelated by a tetrapyrrole ring. It plays a vital role in a wide range of life-essential redox processes, such as detoxification, oxygen transport, regulation of the circadian rhythm, and control of transcription and translation (Feng and Lazar, 2012; Girvan and Munro, 2013; Martínková et al., 2013). Most of the body’s heme is synthesized in terminally differentiating red cells, whose main function is to transport oxygen to all tissues via the heme-containing protein, hemoglobin (Chen and Paw, 2012).

During terminal erythropoiesis, the main function of the erythroid mitochondria is to produce extremely large quantities. Regulation of transcriptional and translational upregulation of genes involved in heme synthesis is fairly well understood. Less is known about how the “housekeeping” mitochondrial homeostatic machinery may regulate the activities of heme synthesis enzymes and mitochondrial iron transport. A few studies have demonstrated that proteins which regulate mitochondrial metabolism also play essential roles in facilitating high rates of heme synthesis in developing erythroid cells (Hildick-Smith et al., 2013; Shah et al., 2012; Sun et al., 2017; Yien et al., 2018). However, the functional interactions between mitochondrial homeostasis and heme synthesis, and the extent to which these regulatory mechanisms are tissue-specific, are not well understood.

We have previously demonstrated that mitochondrial CLPX, a member of the ubiquitious AAA+ (*<u>A</u>TPases <u>a</u>ssociated with various cellular <u>a</u>ctivities*) protein unfoldase family, plays a key role in erythroid differentiation by direct regulation of heme synthesis (Kardon et al., 2015). ClpX functions as a ring-shaped homo-hexamer and is best understood for its function in a proteasome-like enzyme complex with the peptidase ClpP (the ClpXP ATP-dependent protease). In bacteria, ClpX recognizes specific sequences in its protein substrates and unfolds protein tertiary structures by ATP-powered translocation through its central pore, thereby presenting the unfolded polypeptide chain directly to the ClpP proteolytic chamber (Baker and Sauer, 2012). In mammalian cells, CLPX ensures proper folding of mitochondrial respiratory chain proteins (Seo et al., 2016). CLPX therefore regulates the heme synthesis pathway by mediating both activation (Kardon et al., 2015) and degradation (Kubota et al., 2016) of the heme biosynthetic enzymes ALAS1 and ALAS2, which catalyze the committed step of heme synthesis. Collectively, these studies predict that the absence of CLPX will cause accumulation of inactive ALAS protein with a resultant heme deficiency. In this study, we demonstrated that in contrast to previous studies carried out in yeast, vertebrate CLPX is not absolutely required for activation of ALAS2, although it may still activate ALAS2 during terminal erythropoiesis.

To mechanistically dissect the functional role of CLPX in erythroid heme synthesis, we performed loss-of-function studies in murine erythroleukemia (MEL) cells and zebrafish. Our studies demonstrated that the main function of CLPX in erythroid cells was to regulate the turnover of ALAS2. Further, we found that CLPX was required for maximal activity of PPOX and FECH enzymes in the mitochondrial matrix, and may regulate mitochondrial iron utilization during terminal erythroid differentiation. Lastly, we show that in erythroid cells, CLPX does not regulate the function of the mitochondrial respiratory chain protein, SDHB, which was reported to be a CLPXP target (Seo et al., 2016). Collectively, our studies show that CLPX regulates mitochondrial homeostasis and heme synthesis in a cell type specific manner and plays a specialized function in facilitating the high rates of heme synthesis during terminal erythropoiesis.

## Results

CLPX, a ubiquitous mitochondrial protein unfoldase, was recently found to play a central role in heme regulation in eukaryotes via its control of ALAS, the enzyme which catalyzes the committed and rate limiting step of the heme synthesis pathway (Kardon et al., 2015; Kubota et al., 2016; Yien et al., 2017). In *S. cerevisiae*, CLPX facilitates the incorporation of the pyridoxal cofactor (PLP) in ALAS by partially unfolding the ALAS enzyme (Kardon et al., 2015, 2020). The CLPXP protease was also shown to regulate the turnover of ALAS1 in vertebrate cells (Kubota et al., 2016). Taken together, these predict that the absence of CLPX will cause accumulation of inactive ALAS protein with a resultant heme deficiency (Figure 1A). However, a later study in a human patient showed that erythroid cells expressing a CLPX Gly298Asp mutant protein, which disrupted ATPase activity of the CLPX hexamer, had *increased* ALAS2 protein and ALAS2 activity. This led to erythropoietic protoporphyria (EPP) caused by accumulation of protoporphyrin IX, a porphyrin product downstream of ALA (Yien et al., 2017). Collectively, these data suggested that our current understanding of the mechanisms by which CLPX regulates heme synthesis, particularly in vertebrates, is incomplete. In addition, our hypothesis that CLPX deficiency caused anemia due to accumulation of inactive ALAS protein had yet to be directly tested. As the predicted CLPX-interacting regions of yeast ALAS (also known as Hem1) had little similarity to vertebrate ALAS2 (Supplemental Figure 1), we felt that this was an important prediction to test.

**Figure 1.**
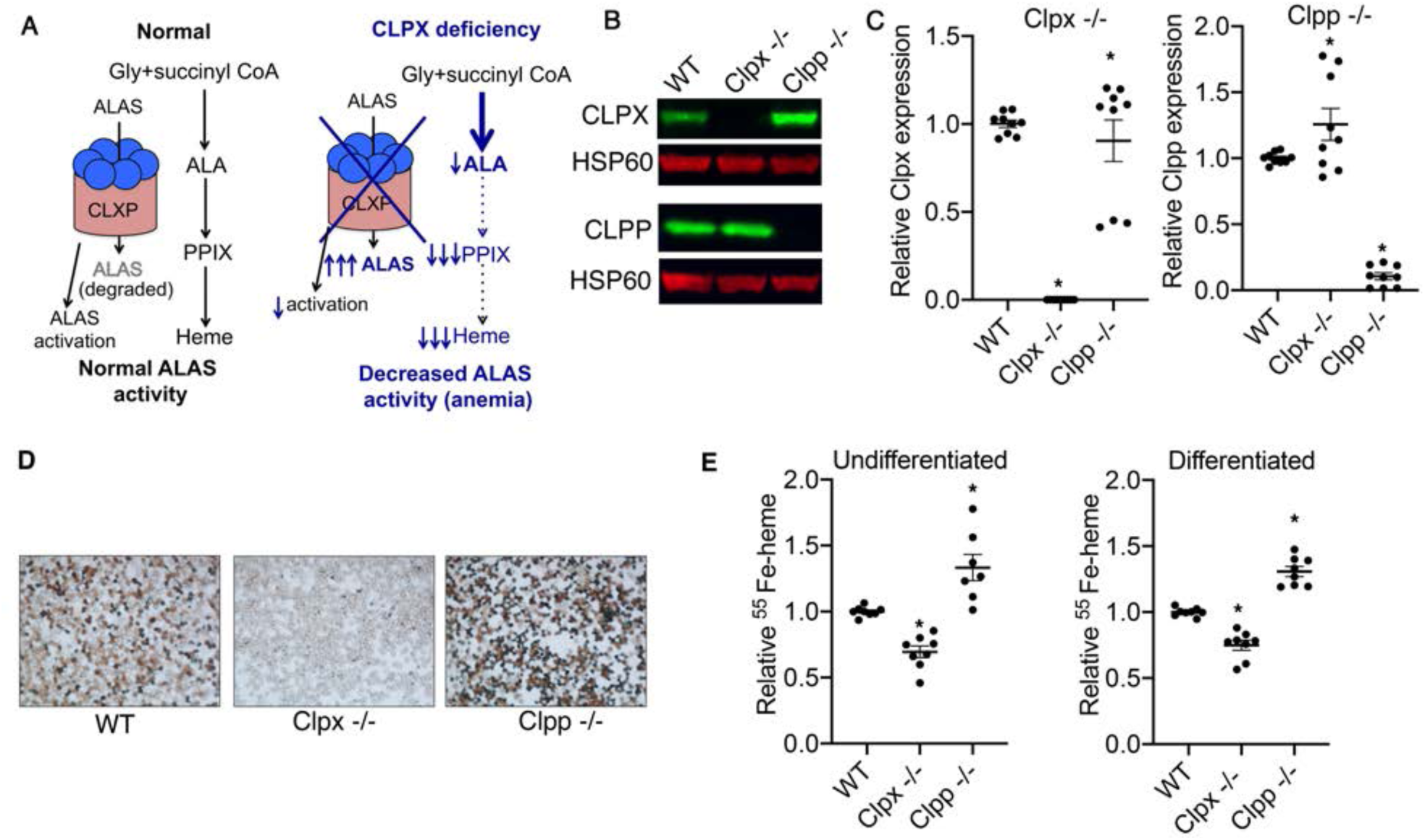
*Clpx -/-* MEL cells have a heme synthetic defect which is distinct of their role in the CLPXP proteolytic complex. (A) Predicted model for the role of the CLPXP protease in heme regulation based on previous work. (Left) In *S*. cerevisiae, CLPX activates the enzyme that catalyzes the rate limiting and committed enzyme of heme synthesis, ALAS. CLPX is also required for erythroid heme synthesis in zebrafish (Kardon et al., 2015). In mammalian cells, the CLPXP proteolytic complex regulates ALAS stability (Kubota et al., 2016; Yien et al., 2017) (Right). Based on these published data, we predict that CLPX-deficient cells will accumulate ALAS protein. This is due to lack of activation by CLPX, and decreased degradation by the CLPXP proteolytic complex. Cells will therefore be heme deficient. In erythroid cells, CLPX deficiency is predicted to result in anemia. (B) Generation of *Clpx -/-* and *Clpp -/-* MEL cells by CRISPR/Cas9. CLPX and CLPP expression in MEL cell lysates were analyzed by western blotting. HSP60 was used as a loading control. (C) Analysis of *Clpx* and *Clpp* mRNA in Clpx -/- and Clpp -/- MEL cells. Gene expression was normalized to expression of *β-actin* mRNA. (n=3, 3 technical replicates) (D) Benzidine staining of *Clpx -/-* and *Clpp -/-* MEL cells that were differentiated in media containing 2% DMSO to analyze heme synthesis during erythroid differentiation. *Clpx -/-* cells were heme deficient, while *Clpp -/-* cells produced more heme than WT cells. (E) Quantitation of *de novo* heme synthesis by ^55^Fe labeling. Both non-differentiating and differentiating *Clpx -/-* cells had a heme synthesis defect. In contrast, *Clpp -/-* cells had increased levels of heme synthesis (n=3, 3 technical replicates). * indicates P<0.05 for student’s t test for statistically significant differences between WT and mutant cell lines.

To test our hypothesis, we used CRISPR/Cas9 to knock out the *Clpx* and *Clpp* genes in mouse erythroleukemia (MEL) cells. Western blot and qRT-PCR analysis of the cell lines indicated that the *Clpx -/-* and *Clpp* -/- cells produced almost no CLPX or CLPP protein (Figure 1B) and mRNA (Figure 1B and 1C). Differentiation of *Clpx -/-* and *Clpp* -/- MEL cells, followed by benzidine staining for heme, indicated that *Clpx -/-* cells were heme deficient, as would be predicted from previously published observations that *clpx* zebrafish morphants were anemic (Figure 1A) (Kardon et al., 2015). This was not caused by defects in mitochondrial biogenesis or cell death, which we assayed by mitotracker red staining (Supplemental Figure 2). Consistent with our hypothesis, *Clpp -/-* cells, which are predicted to have defective CLPXP-catalyzed degradation and therefore accumulate active ALAS2 and increased heme synthesis, had increased heme content (Figure 1D). To quantify the effects of *Clpx* and *Clpp* on erythroid heme synthesis, we labeled newly synthesized heme with ^55^Fe. We extracted heme from these cells with cyclohexanone and quantitated ^55^Fe-heme by liquid scintillation. Both undifferentiated and differentiated *Clpx -/-* MEL cells synthesized decreased levels of heme. In contrast, *Clpp -/-* MEL cells synthesized increased quantities of heme (Figure 1E). These data confirm previous observations that CLPX is required for heme synthesis, while CLPP is not, but may play a role in modulating heme synthesis. Further, these data also confirm that in the heme synthesis pathway, CLPX carries out functions that are independent of its role in the CLPXP proteolytic complex.

**Figure 2.**
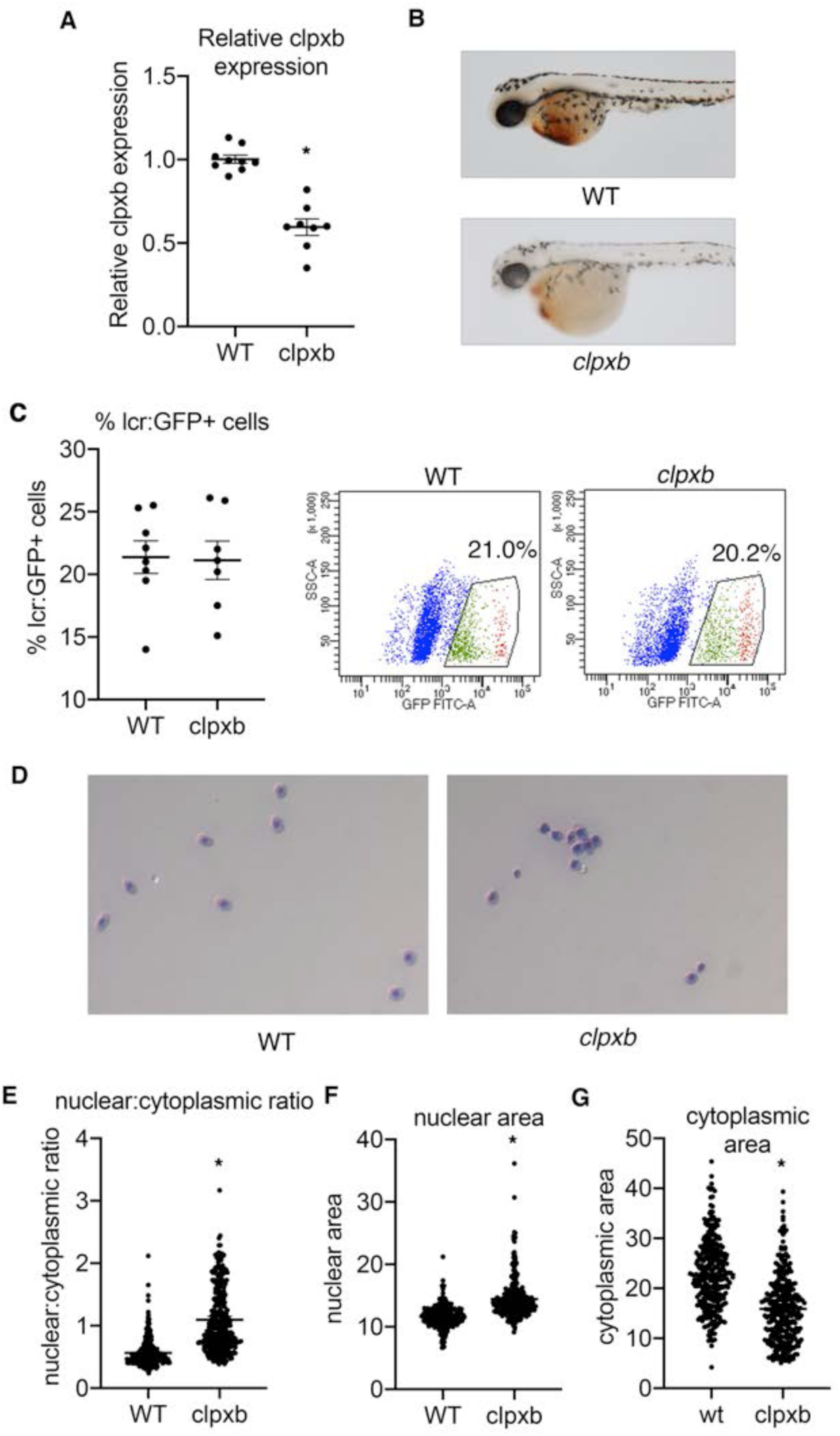
*clpxb* mutant zebrafish embryos develop normal numbers of red cells, but are heme deficient and have dysregulated red cell morphology. (A) *clpxb* mRNA levels are decreased by approximately 50% in mutant embryos. (B) *clpxb* is required for erythroid hemoglobinization and other functions in zebrafish development. *clpxb* mutant embryos are anemic at 48 hpf and are dead by 6 dpf. (C) *clpxb* is not required for erythroid differentiation. *clpxb/clpxb*; Tg(*lcr:GFP*) zebrafish embryos had similar percentages of GFP^+^ erythroid cells to control Tg(*lcr:GFP*) embryos. This indicates that *clpxb* is not required for erythroid specification. (D) Erythroid cells from 72hpf WT and *clpxb* mutant zebrafish embryos Tg(*lcr:GFP*) embryos were sorted onto a slide, giemsa stained and imaged. Erythroid cells from *clpxb* mutant embryos were more variable in size and appearance, had smaller cytoplasmic areas, and some cells had larger nuclei. 298 erythroid cells were quantitated from WT and *clpxb* Tg(*lcr:GFP*) embryos. (E) The nuclear to cytoplasmic ratio of WT and *clpxb* erythroid cells was quantitated, with *clpxb* erythroid cells having higher nuclear: ratios. In addition, the nuclear: cytoplasmic ratios of *clpxb* erythroid cells exhibited higher variability (F test: P< 0.05). (F) The increased nuclear:cytoplasmic ratio is caused by both increased nuclear area in *clpxb* erythroid cells and their (G) decreased cytoplasmic area. * indicates P<0.05, student’s t test.

To further understand the role of *Clpx* in erythropoiesis *in vivo*, we characterized a zebrafish *clpxb* mutant (*clpxb*^*sa38141*^) that was obtained from Zebrafish International Resource Center (ZIRC). The mutant carries a E129X nonsense mutation in its *clpxb* gene, which removes a large portion of the protein, including substantial sections of the large AAA+ domain, and completely removes the small AAA+ domain, which are required for ATP binding. Homozygous mutant fish embryos express about half the amount of *clpxb* mRNA as wild-type zebrafish, due to nonsense mediated degradation (Figure 2A). At 48 hpf, *clpxb* mutant embryos were anemic, consistent with previous observations (Kardon et al., 2015) (Figure 2B). To determine if the anemia was caused by defects in erythroid differentiation, we crossed *clpxb* embryos to *Tg(globin-lcr:GFP)* zebrafish in erythroid cells are marked by GFP (Ganis et al., 2012) and quantitated erythroid cells by flow cytometry. In contrast to *clpxa* morphants, *clpxb* mutant zebrafish did not exhibit a decrease in erythroid cell numbers, indicating that *clpxb* is not required for erythroid specification (Figure 2C). To further characterize the erythroid cells in *clpxb* mutant zebrafish, we sorted GFP^+^ cells and Giemsa stained them (Figure 2D). We observed that in contrast to wild-type erythroid cells, the *clpxb* mutant cells had variable cell size, larger and more variable nuclear:cytoplasmic ratios (Figure 2E) that were caused by increases in nuclear size (Figure 2F) and decreases in cytoplasmic area (Figure 2G). The cells also had ruffled membranes and irregular morphology, in contrast to the smooth, oval shaped appearance of the wild-type cells and nuclei (Figure 2D, E). These data indicate that *clpxb* is required for erythroid heme synthesis and the morphological processes of terminal erythropoiesis, but is not required for erythroid lineage determination or regulation of erythroid cell number.

To determine if CLPX regulates ALAS2 protein stability, we first analyzed the expression of steady state mRNA and protein. In undifferentiated MEL cells, qPCR analysis revealed that *Alas2* mRNA expression *decreased* in *Clpx -/-* and *Clpp -/-* cells, relative to wild-type cells. In differentiated MEL cells, ALAS2 mRNA decreased in *Clpx -/-* cells, relative to wild-type cells, and remained unchanged in *Clpp -/-* cells (Figure 3A(i)). ALAS2 steady state protein expression increased in both *Clpx -/-* and *Clpp -/-* cells (Figure 3A (ii)). The increase in ALAS2 protein levels in *Clpx -/-* and *Clpp -/-* cells, without an increase in *Alas2* mRNA levels suggests that CLPX and CLPP may regulate ALAS2 protein stability, similar to its role in ALAS1 (Kubota et al., 2016). In contrast to previous observations in prostate adenocarcinoma cells, we did not observe increases in SDHB protein, or significant changes in SDHB activity in *Clpx -/-* and *Clpp -/-* cells (Supplemental Figure 3).

**Figure 3.**
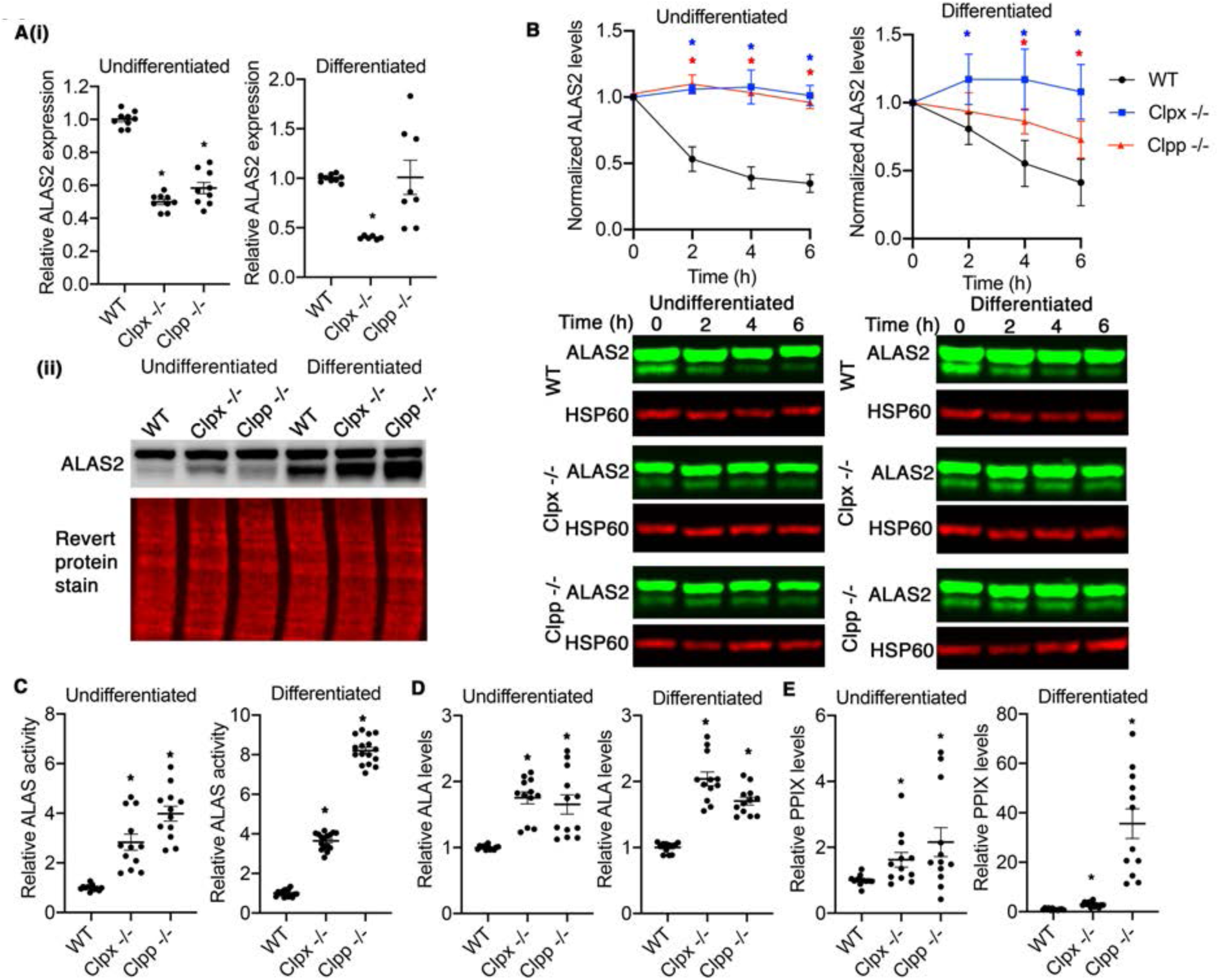
CLPX regulates protein stability of ALAS2, but is not required for ALAS2 activity. (A) (i) ALAS2 mRNA expression is decreased or unaltered in the absence of CLPX and CLPP, (ii) but its protein levels are increased in *Clpx -/-* and *Clpp -/-* cells. (B) ALAS2 stability is increased in both *Clpx -/-* and *Clpp -/-* cells. (C) In contrast to predictions that CLPX is required for ALAS activity, ALAS2 activity was significantly increased in *Clpx -/-* cells. This increase in activity was also observed in *Clpp -/-* cells. While the increase in ALAS2 activity was observed in *Clpx -/-* and *Clpp -/-* cells, the increase in ALAS2 activity in *Clpp -/-* cells was far more pronounced in differentiating MEL cells. (D) Consistent with the increase in ALAS2 activity, *Clpx -/-* and *Clpp -/-* cells produced significantly more ALA (almost 2x) than WT cells. (E) *Clpx -/-* and *Clpp -/-* cells produced significantly more PPIX than WT cells. The increase in PPIX was similar to the increase of the ALA precursor in undifferentiated cells. The increase in PPIX in differentiated cells exceeded the increase in ALA production by several fold, in *Clpx -/-* cells and dramatically more so in *Clpp -/-* cells.

To determine if CLPXP regulates ALAS2 stability, we monitored the rate of ALAS2 turnover in WT, *Clpx -/-* and *Clpp -/-* cells following inhibition of protein translation by cycloheximide (Laghmani et al., 2016; Quadrini and Bieker, 2006; Schneider-Poetsch et al., 2010; Yien and Bieker, 2012; Yien et al., 2017). Over 6 hours, we monitored the levels of ALAS2 protein by Western blot analysis. ALAS2 levels were normalized to the levels of HSP60, a mitochondrial resident protein whose expression did not appreciably change during cycloheximide treatment. We observed that in both differentiating and non-differentiating cells, the absence of *Clpx -/-* and *Clpp -/-* stabilized ALAS2 protein (Figure 3B). These results were consistent with observations that CLPXP regulates ALAS1 turnover (Kubota et al., 2016).

To test the hypothesis that CLPX regulates heme synthesis by activating ALAS in vertebrate cells, we carried out ALAS enzyme activity assays in WT, *Clpx -/-* and *Clpp -/-* cells. To our surprise, both undifferentiated and differentiated *Clpx -/-* cells exhibited significantly increased ALAS activity, indicating that CLPX is not absolutely required for ALAS activation. Strikingly, *Clpp -/-* cells had even higher ALAS enzyme activity than *Clpx -/-* cells. This dramatic increase in ALAS activity was most pronounced in differentiated *Clpp -/-* cells (Figure 3C). The relative increase of ALAS activity in *Clpp -/-* relative to *Clpx -/-* cells cannot be attributed to changes in protein expression or stability (Figures 3A and B) suggesting that CLPP may play an inhibitory role in ALAS activation. An alternative explanation is that presence of CLPX, which may activate ALAS, combined with the absence of CLXP-mediated degradation, synergized to superactivate ALAS. We quantitated ALA, the product of ALAS to determine if these differences of ALAS activity affected synthesis of its product and downstream heme synthesis. While the absence of CLPX or CLPP caused an increase in ALA levels, both *Clpx -/-* and *Clpp -/-* cells had similar ALA levels (Figure 3D). Finally, we quantitated protoporphyrin IX (PPIX), the terminal heme intermediate, to determine if porphyrin levels mirrored changes in ALA synthesis. First, we observed that in both differentiated and undifferentiated cells, *Clpx -/-* and *Clpp -/-* cells had increased PPIX levels. However, to our surprise, while *Clpx -/-* and *Clpp -/-* cells had *similar* increases in ALA levels, *Clpp -/-* cells contained significantly more PPIX than *Clpx -/-* cells. This was especially apparent in differentiated cells, suggesting that in erythroid cells, CLPXP may regulate heme synthesis downstream of ALA production (Figure 3E). Collectively, the results in Figure 3 demonstrate that contrary to earlier predictions (Kardon et al., 2015), the heme defect in *Clpx -/-* erythroid cells was not a result of a decrease in ALAS activation, nor a defect in porphyrin production.

Since CLPX is situated in the mitochondrial matrix (Rhee et al., 2013), we hypothesized that CLPX may regulate terminal processes in the heme synthesis pathway that occur within the mitochondrial matrix, such as PPOX or FECH activity (Rhee et al., 2013) or mitochondrial iron metabolism (Shaw et al., 2006). We carried out Western blot analysis of PPOX and FECH protein levels to determine the effect of CLPXP on their expression. Although PPOX levels remained largely similar in WT, *Clpx -/-* and *Clpp -/-* cells, FECH levels were decreased (Figure 4A). *Clpx -/-* cells experienced a decrease in PPOX activity relative to either WT or *Clpp -/-* cells (Figure 4B). FECH activity was decreased in *Clpx -/-* cells relative to wild-type and *Clpp -/-* cells in non-differentiating cells. However, in *differentiating* cells, both *Clpx -/-* and *Clpp -/-* cells had decreased FECH activity. (Figure 4C). As *Clpp -/-* cells did not exhibit a heme synthesis defect, it was unlikely that the FECH defect was the sole cause of the heme defect observed in *Clpx -/-* cells.

**Figure 4.**
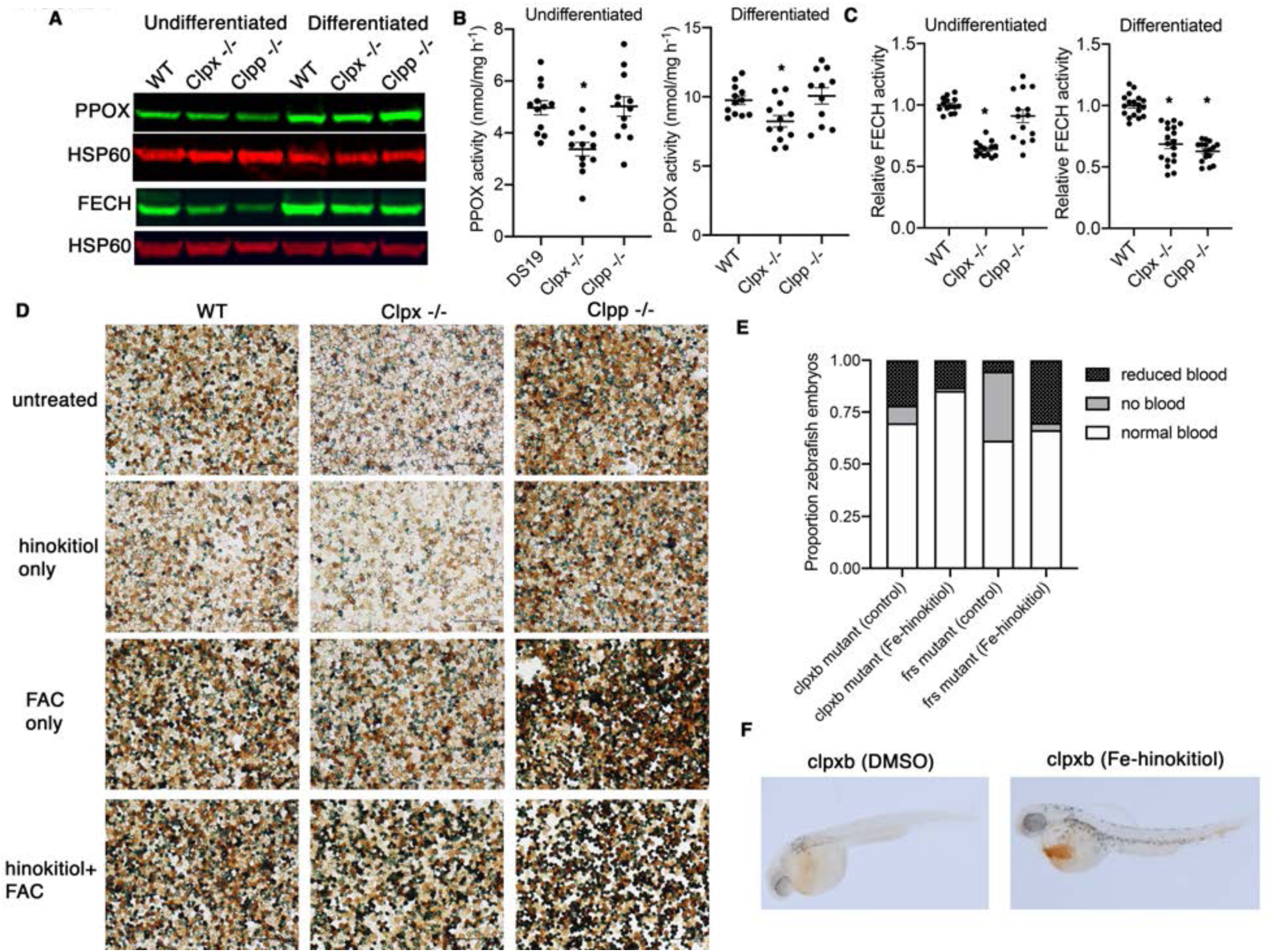
CLPX regulates mitochondrial iron metabolism in erythroid cells. (A) PPOX protein levels remain unaltered in *Clpx -/-* and *Clpp -/-* cells. However, FECH protein levels are decreased. This may account for the decrease in FECH activity. (B) PPOX activity is decreased in *Clpx -/-* cells, in both undifferentiated and differentiated cells. PPOX activity is unaltered in *Clpp -/-* cells. (C) FECH activity is decreased in undifferentiated *Clpx -/-* cells, relative to wild-type cells. The FECH activity of *Clpp -/-* cells were unaltered. In differentiating MEL cells, FECH activity is decreased in both *Clpx -/-* and *Clpp -/-* cells. This decrease may partially account for the increase in PPIX levels, relative to ALA levels. (D) Supplementation of *Clpx -/-* cells with iron, in the form of ferric ammonium citrate (FAC), and in combination with hinokitiol (a lipophilic iron chelator) restores heme synthesis in differentiating MEL cells to near wild-type levels. *Clpp -/-* cells synthesized greater quantities of heme than wild-type cells under untreated conditions, and during iron supplementation with either FAC alone or in combination with hinokitiol. (E) Addition of iron-hinokitiol to *clpxb* homozygous mutant 48 hpf zebrafish embryos significantly reduced the numbers if anemic fish. As a control, we treated *frs* homozygous mutant zebrafish embryos with iron-hinokitiol. Iron-hinokitiol significantly decreased the number of bloodless fish and ameliorated the anemia. (F) Representative images of *clpxb* zebrafish that were treated with Fe-hinokitiol. Hemoglobinization of erythroid cells was significantly increased.

We also considered the possibility that CLPX also regulated mitochondrial iron metabolism in erythroid cells. To determine if this was so, we attempted to chemically complement the heme defect in *Clpx -/-* cells with iron, in the form of ferric ammonium citrate (FAC), in combination with a lipophilic iron chelator, hinokitiol, that can transport iron across a concentration gradient into iron-deficient sub-cellular compartments (Grillo et al., 2017). In untreated cells, *Clpx -/-* cells exhibited less heme staining than WT or *Clpp -/-* cells. This was also the case with the cells were treated with hinokitiol without supplemental iron. Treatment with FAC increased heme content in all 3 cell types. Notably, addition of hinokitiol in combination with FAC further increased heme content in *Clpx -/-* cells to levels close to wild-type. These results suggested that a mitochondrial iron defect could contribute to the heme defect unique to *Clpx -/-* cells (Figure 4D). We repeated this experiment in zebrafish, by treating clutches of embryos spawned from *frs* or *clpxb* heterozygous incrosses with either vehicle (DMSO) or iron-hinokitiol. *frs* mutant zebrafish, which lack expression of *mfrn1*, the mitochondrial iron transporter (Christenson et al., 2018; Shaw et al., 2006), served as a positive control for this experiment as their anemia shown to be ameliorated by iron-hinokitiol (Grillo et al., 2016). As expected, the numbers of severely zebrafish embryos (almost no heme staining) in *frs* incrosses were sharply reduced when treated with iron-hinokitiol. In comparison, treatment of embryos from *clpxb* heterozygote incrosses with iron-hinokitiol significantly decreased numbers of embryos in both “severely anemic” and “anemic” categories, but the results were not as clear cut as the *frs* mutant zebrafish, which have a clear iron defect (Figure 4E). These intermediate results indicate that iron deficiency may play a role in anemia in the *clpxb* mutant embryos (Figure 4F), but was not a sole cause of the anemia. Western blot analysis indicates that neither *Clpx -/-* or *Clpp -/-* cells had a defect in MFRN1 expression (Supplemental Figure 4).

Collectively, our results indicate that the primary mechanism by which CLPX regulates ALAS in vertebrate cells is by control of its turnover, rather than its activity. We have also shown that CLPX is required for maximal PPOX and FECH activity and may play a role in regulating and mitochondrial iron metabolism in differentiating erythroid cells. The porphyrin accumulation observed in these cells is augmented by the increase in ALAS2 activity in these cells, caused by an increase in protein levels and stability (Supplemental Figure 5).

## Discussion

Our results demonstrate that although *Clpx -/-* erythroid cells are heme deficient, like *Δmcx1* yeast, *Clpx -/-* erythroid cells have elevated ALA content and ALAS activity, unlike in *Δmcx1* yeast. As PLP is required for ALAS enzyme activity (Hunter and Ferreira, 2011), these data show that vertebrate ALAS2 does not absolutely require CLPX to facilitate PLP incorporation into the enzyme complex. However, ALAS activity in *Clpp -/-* cells is more dramatically elevated than that of *Clpx -/-* cells even though the steady state protein levels in *Clpx -/-* and *Clpp -/-* cells are similar (Figure 3A). This suggests that CLPX may still play a nonessential role in ALAS activation, leading to *Clpp -/-* cells exhibiting a “superactivated” ALAS phenotype as CLPX activates the accumulated ALAS2. It was evident that the heme defect in vertebrate *Clpx -/-* cells was not caused by a deficiency in ALAS activity or ALA production. It is also evident as PPIX production was elevated in both *Clpx -/-* and *Clpp -/-* cells, which was likely caused by a combination of elevated ALA, and downstream heme synthesis pathway and iron metabolism defects. Non-differentiating *Clpx -/-* cells exhibited decreased PPOX and FECH activity, likely accounting for the decrease in heme synthesis. The elevation of PPIX levels in these cells is likely due to a combination of increased protoporphyrinogen IX (PPgenIX) caused by the combination of ALA overproduction and PPOX deficiency. PPgenIX auto-oxidises to PPIX in cells when lysed for HPLC analysis, leading to increased measurable PPIX even with decreases in PPOX activity in *Clpx -/-* cells. In differentiating MEL cells, we observed that FECH activity was decreased in *Clpp -/-* cells, compounding the PPIX accumulation, although this did not appear to cause a heme deficiency in these cells. The heme defect in *Clpx -/-* cells was even further exacerbated by an iron metabolic defect, which was ameliorated by addition of exogenous iron (Figure 4D-F). Lastly, levels of MFRN1, the mitochondrial iron importer, were similar in all three cell lines. We concluded that the specific heme defect in *Clpx -/-* terminally differentiating mouse and zebrafish erythroid cells and zebrafish was primarily caused by a decrease in PPOX activity, which limited the amount of PPIX formed. The heme defect was exacerbated by a defect in FECH activity and mitochondrial iron metabolism. This is supported by the partial rescue of erythroid heme synthesis by exogenous iron (Figure 4D-F).

The CLPXP proteolytic complex, which has been extensively studied in bacteria, consists of an ATP-dependent CLPX subunit which unfolds and translocates protein substrates into a CLPP proteolytic barrel, which then degrades unfolded protein substrates (Olivares et al., 2016). Although this structure is conserved in vertebrates, there are several reports that CLPX has roles that are distinct from its function in CLPXP mediated proteolysis (Cheong et al., 2020; Kardon et al., 2015; Seo et al., 2016; Wang et al., 2018). This is most strikingly illustrated by the fact that *Clpx -/-* mouse embryos die by E8.5 with severe gastrulation defects (Cheong et al., 2020), while *Clpp -/-* mice survive to adulthood (Wang et al., 2018). Our data adds to studies that demonstrate that CLPX has roles that are independent of its function within the CLPXP proteolytic complex. Our work shows that PPOX protein levels remained unaltered in *Clpx -/-* and *Clpp -/-* MEL cells, while PPOX activity was decreased in *Clpx -/-* cells, suggesting that CLPX plays a specific role in PPOX activation. Further, our iron complementation studies suggest that *Clpx -/-*, but not *Clpp -/-* erythroid cells have a mitochondrial iron defect. Collectively, our studies suggest that CLPX is required for regulating the activities and stabilities of mitochondrial matrix proteins that are required for optimal heme synthesis in erythroid cells.

The functional interactions between CLPX/P and the erythroid heme synthesis pathway that are described in this manuscript are highly complex and are likely to be tissue and developmental stage specific. The regulation of mitochondrial energy metabolism and physiology by CLPX is an emerging theme in the field (Fischer et al., 2015; Seo et al., 2016). Of note, several CLPXP binding proteins that are predicted to be substrates are iron-sulphur cluster proteins that are involved in oxidative phosphorylation or iron-sulphur cluster assembly (Fischer et al., 2015). In a mouse model of Freidreich ataxia in which frataxin was specifically deleted in striated muscles, there was an increase in expression of the CLPP and LON proteases that was concomitant with a decrease in mitochondrial Fe-S proteins. This included FECH, NDUFS3 and SDHB proteins, which are involved in heme synthesis and oxidative phosphorylation (Guillon et al., 2009). These data contrast with the decrease in FECH protein and activity that we observed in our *Clpx -/-* and *Clpp -/-* erythroid cell lines (Figure 4A, C). We also observed that unlike previous studies in prostate adenocarcinoma cells (Seo et al., 2016), SDHB protein levels and activity were relatively unchanged in *Clpx -/-* and *Clpp -/-* cells (Supplemental Figure 3A). While there is convincing data to show that CLPX/P plays an important role in the regulation of cellular metabolism, particularly in pathways that require iron, it is likely that the specific roles of CLPXP in cell metabolism are regulated by cell-specific binding proteins, as individual cell types have specific metabolic needs. For instance, erythroid cells shift from the use of the Krebs’ cycle and oxidative phosphorylation to exclusive dependence on glycolysis as they terminally differentiate mitochondria (Gasko and Danon, 1972c, 1972a, 1972b). During terminal differentiation, the main function of erythroid mitochondria shifts towards heme production for hemoglobin synthesis (Chen et al., 2013; Chung et al., 2017; Yien et al., 2014, 2018; Zhang et al., 2003). It is therefore unsurprising, but revealing, that CLPXP regulates erythroid heme synthesis at multiple points in the pathway, but does not seem to be as essential for regulating oxidative phosphorylation proteins as it does in other cell types.

The data described in this manuscript also shed light on how by a dominant *CLPX*^*G298D*^ gain-of-function heterozygous mutation caused erythroid protoporphyria (Yien et al., 2017). Our previous manuscript showed that the mutant protein lacked ATPase activity and hetero-oligomerized with wild-type CLPX, decreasing the ATPase activity of the CLPX hexamer. Based on previously published data, we proposed that the WT/G298D hetero-hexamer possessed sufficient ATPase activity to activate ALAS, but had insufficient ATPase activity to translocate ALAS into the CLPP proteolytic barrel, causing stabilization of active ALAS and subsequent overproduction of ALA and PPIX, leading to EPP symptoms. Our current data indicate that CLPX is not required for activation of mammalian ALAS, suggesting that CLPX-EPP could be caused by stabilization of ALAS alone. However, our data also suggest that CLPX/P is required for optimal FECH activity, and loss of CLPX/P can cause a decrease of FECH activity. This decrease in FECH activity can further exacerbate PPIX accumulation caused by ALA overproduction and may be compounded by an iron defect.

Given the involvement of CLPX and CLPXP in essential mitochondrial processes such as respiration and heme synthesis, it is now known that CLPX mutations contribute to metabolic disorders in both human patients and animal models (Gispert et al., 2013; Guillon et al., 2009; Jenkinson et al., 2013; Yien et al., 2017) Our biochemical studies in erythroid cells indicate that CLPX is required for optimal heme synthesis in erythroid cells by control of mitochondrial iron utilization. The heterogeneity of CLPX-deficient phenotypes indicates that CLPX is likely to regulate mitochondrial metabolism in a cell-specific manner. Based on our studies, we speculate that the CLPX hexamer by itself, and as part of the CLPXP protease, interacts with cell-specific factors and play an important role in coupling mitochondrial metabolism with cellular requirements in a tissue-type and developmental specific manner. Our studies reveal that CLPXP is critical for correct regulation of erythroid heme synthesis, particularly during terminal maturation. Unraveling the complexities of CLPX function in metabolism will be key to understanding how mitochondrial function is regulated in specific cell types under different conditions, and will reveal therapeutic strategies to treat metabolic diseases and mitochondriopathies in a targeted manner.

## Materials and Methods

### Vertebrate animal study approval

All vertebrate animal studies were performed in compliance with Institutional Animal Care and Use Committee protocols at the University of Delaware.

### Cell and zebrafish lines

MEL DS19 cells were obtained from Arthur Skoultchi (Albert Einstein College of Medicine, New York). The *clpxb*^*sa38141*^ zebrafish mutant line was obtained from ZIRC. Zebrafish lines were genotyped by amplifying the region around the *clpxb* mutation with the following primers: Fwd 5’-ACTGGAGAAGAGCAACATTGTGTTG −3’ and Rev 5’-CGATGTCTTCGCCCACATATCC −3’. The mutation was confirmed by Sanger sequencing.

### Generation of CRISPR cell lines

CRISPR guide sequences were designed to direct two cleavages at the target gene loci to generate a chromosomal deletion (Canver et al., 2014; Cong et al., 2013). CRISPR guide sequences were designed to have a unique 12 bp seed sequence 5’-NNNNNNNNNNNN-NGG-3’ in the mouse genome (http://www.genome-engineering.org) to minimize off-target cleavages. We targeted exons 3 and 11 of the *Clpx* gene with the following sequences: 5’CACCGAAGGAAGCAGTAAGAAATCG 3’ (exon 3) and 5’ CACCGGATCTTGCTAACCGAAGTGG 3’ (exon 11). We targeted exons 1 and 4 of the *Clpp* gene with the following sequences: 5’ CACCGAGGCCCGGGTGGCTGTGGAG 3’ (exon 1) and 5’ CACCGTTGGACAGGCTGCCAGCATG 3’ (exon 4). CRISPR guides were cloned into pX330 plasmid (Addgene) with BbsI ligation as previously described (Ran et al., 2013).

CRISPR/Cas9 constructs were delivered to mouse erythroleukemia (MEL) cells by electroporation using a Biorad electroporator. CRISPR/Cas9 constructs were co-electroporated with pEF1*α* at a 9:1 ratio. Cell selection was carried out as previously described (Yien et al., 2018). Clones were screened for loss of CLPX or CLPP protein by western analysis and qRT PCR.

### Functional complementation assays with iron in cultured cells

MEL cells were differentiated with 2% DMSO for 72 h. For functional complementation assays with hinokitiol, cells were concurrently treated with 10 μM ferric ammonium citrate and hinokitiol (1 μM). Zebrafish embryos were treated with 1μM hinokitiol and 10 μM ferric citrate (Grillo et al., 2017).

### Iron radiolabeling and radio iron-heme measurements

^55^FeCl_3_ (Perkin Elmer) was loaded onto transferrin as previously described (Roy et al., 1999). Metabolic labeling with ^55^Fe-transferrin and quantitation of ^55^Fe-heme were carried out as previously described (Yien et al., 2014).

### Western Analysis

The following primary antibodies and dilutions were used in this study: ALAS2 (Abcam, ab184964), 1:1000; CLPX (Abcam, ab168338), 1:1000; CLPP (Abcam, ab12482), 1:1000; FECH (EMD Millipor Corp, ABS2124), 1:500; PPOX (Abnova, H00005498-M01), 1:500; HSP60 (Abcam ab59457) 1:750; HSP60 (Sigma, A5316), 1:750; *β*-actin (Santa Cruz, sc47778), 1:1000; SDHB (Abcam ab14714), 1:1000; MFRN1 (Proteintech, 26469-1-AP), 1:500. Fluorescent secondary antibodies were obtained from Li-COR and probed at a dilution of 1:10,000. Secondary antibodies for chemiluminescence were from Sigma and used at a dilution of 1:10,000.

### qRT-PCR

qRT-PCR probes were obtained from ThermoFisher Scientific. For quantitation of zebrafish *clpxb* mRNA, we custom. designed a probe (5’ CTGGGACCCACTGGATCAGGTAAAACTCTCCTGGCCCAAACA 3’) which bound between exons 8-9. Actin was used as a loading control (Dr03432610_m1). For mouse cell lines, the following probes were used: *Alas2* Mm00802083_m1; *Clpx* Mm00488586_m1; *Clpp* Mm00489940_m1; *β*-actin Mm02619580_g1.

### CHX Block

Inhibition of translation was carried out with 100 μg/mL CHX (Sigma-Aldrich) as previously described (Yien and Bieker, 2012; Yien et al., 2018). Cells were harvested at indicated times after CHX treatment. Proteins whole cell lysates were resolved by SDS/PAGE. ALAS2 and HSP60 were detected by Western blot analysis, and protein levels were quantified using a LI-COR imager and normalized to fluorescence at t=0h.

### Single embryo cell extraction and FACS sorting of GFP^+^ erythroid cells

Embryos were anesthetized tricaine and individual embryos were place on ice prior to processing and materials and solutions were chilled to maintain integrity. Embryos were rinsed with PBS, crushed thoroughly with a tube pestle, and remaining tissue mechanically dissociated. All remaining steps were done on ice. The cell solution was filtered, rinsed with PBS. Cells were collected by centrifugation and resuspended in 0.05% gluteraldehyde fixative. Fixed cells were collected by centrifugation and rinsed with PBS and PBS + 0.1%BSA. Cells were sorted using a FACSaria Fusion at 600 cells per slide. Remaining cells were spun down for genotyping.

### Wright-Geimsa Staining

Slides were fixed in Methanol. Slides were then rinsed in water, dried, and Wright-Geisma stained using recommended conditions. Slides were rinsed in water, air dried, and cover slipped using cytoseal 60.

### Biochemical Analyses

Activity assays for ALAS, FECH and PPOX enzymes, and sample ALA and PPIX levels were measured at the Iron and Heme Core of the Center for Iron and Heme Disorders, University of Utah. ALAS activity and ALA content were measured as described (Bergonia et al., 2015). The FECH assay method was based mainly on a method as described (Rossi et al., 1988). PPIX was extracted from the samples using the method described by (Peter et al., 1978). The PPOX assay protocol was modified from (Li et al., 1987). More details of the methods are posted at http://cihd.cores.utah.edu/ironheme/#1467136333172-56f37fbb-2816.

## Acknowledgements

This work was supported by the Cooley’s Anemia Foundation (to Y.Y.Y) and the National Institutes of Health grants K01 DK106156, R03 DK118307, P01 HL032262, NIH/NIGMS P20GM104316, R35 GM1133560 and a U. Utah CIHD pilot grant (Y.Y.Y.), and U54 DK083909 (J.D.P).

