## Supplemental data for "CLPX regulates erythroid heme synthesis by control of mitochondrial heme synthesis enzymes and iron utilization"

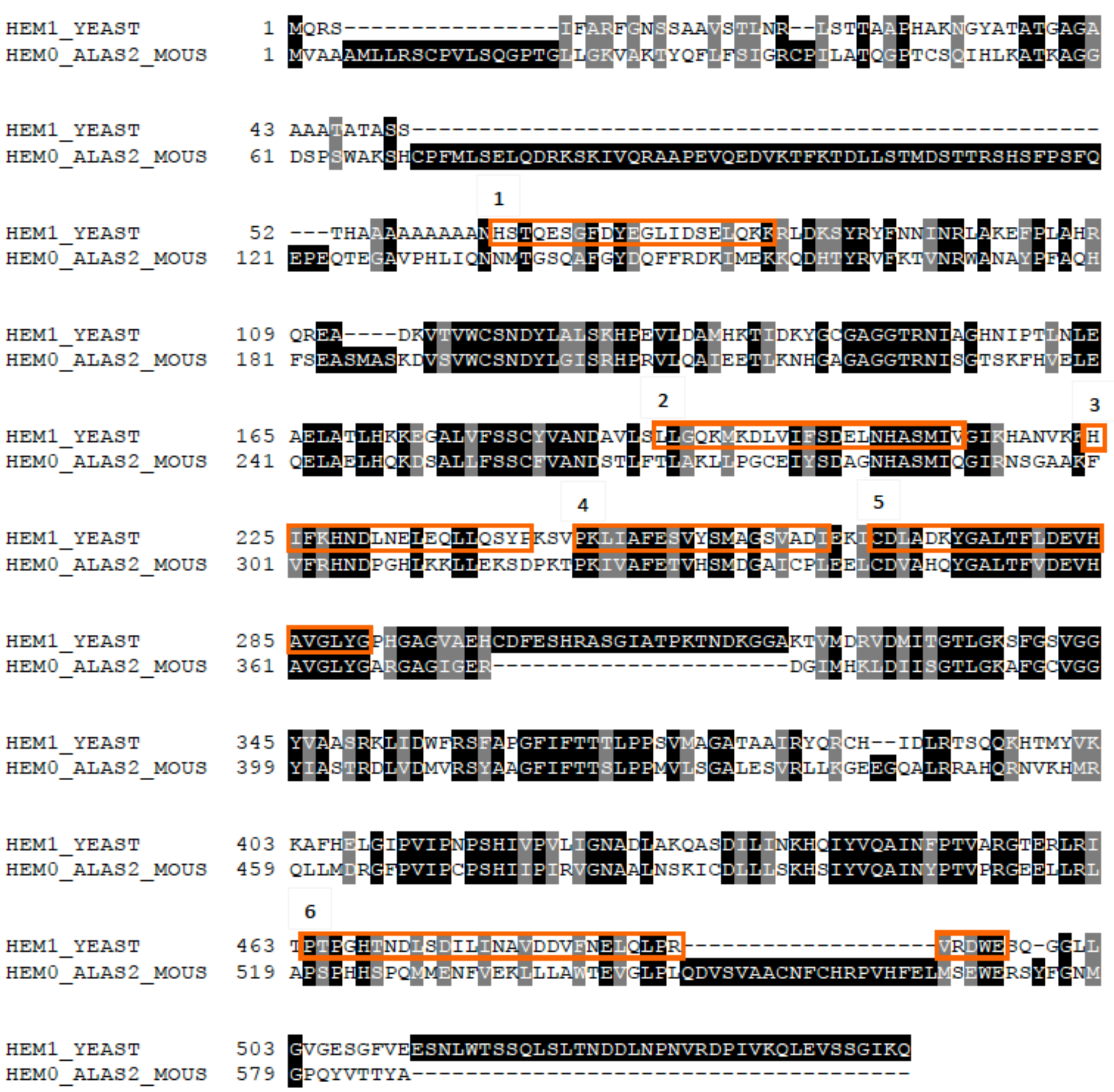


**Supplemental Figure 1.** Alignment of *S. cerevisiae* Hem1 with mouse ALAS2. The boxed regions on yeast Hem1 are sequences that bound to Mcx1 (Clpx) (Kardon, Moroco, Engen, & Baker, 2020).

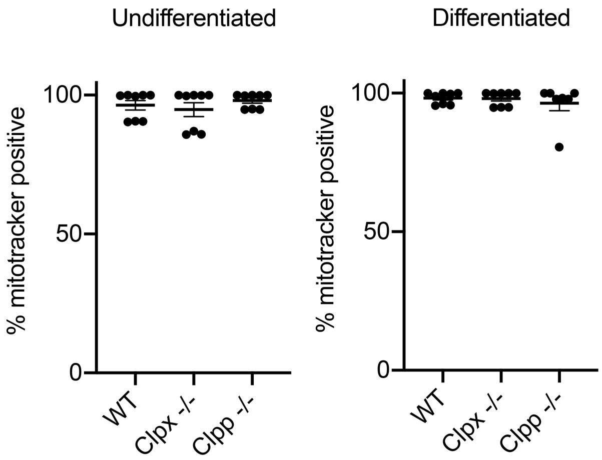


**Supplemental Figure 2. Mitotracker staining of WT, *Clpx -/-* and *Clpp -/-* MEL cells.**

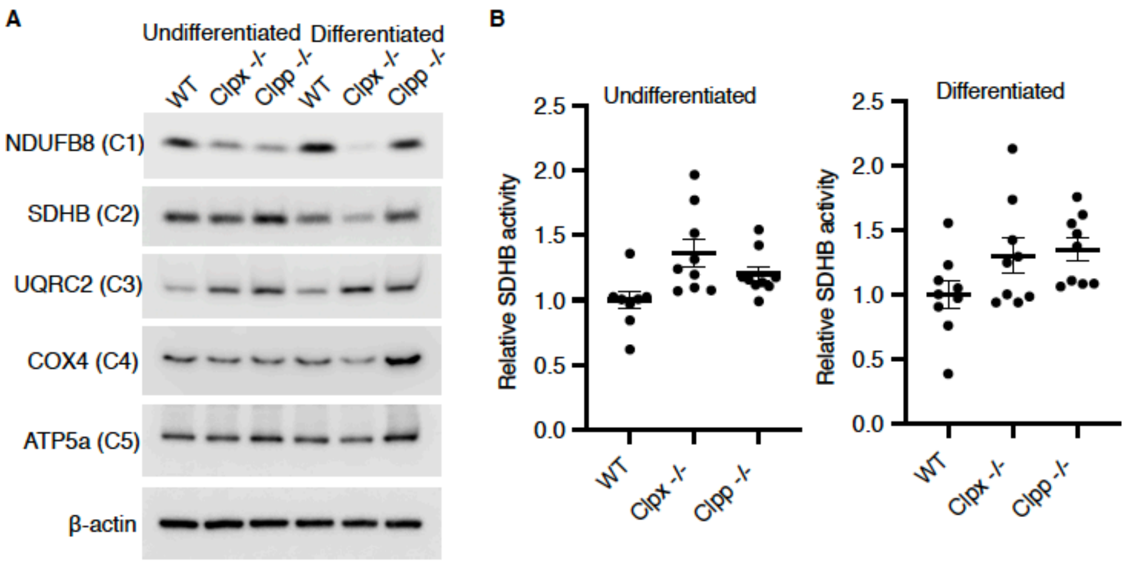


**Supplemental Figure 3. SDHB protein levels and activity are unaltered in *Clpx -/-* and *Clpp -/-* MEL cells.** (A) Western blot of oxidative phosphorylation complex proteins. (B) SDHB activity assays in undifferentiated (left) and differentiated (right) MEL cells.

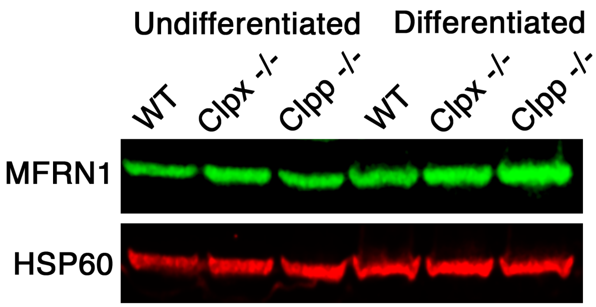


**Supplemental Figure 4. Western blot analysis of MFRN1 levels in *Clpx -/-* and *Clpp -/-* MEL cells.**

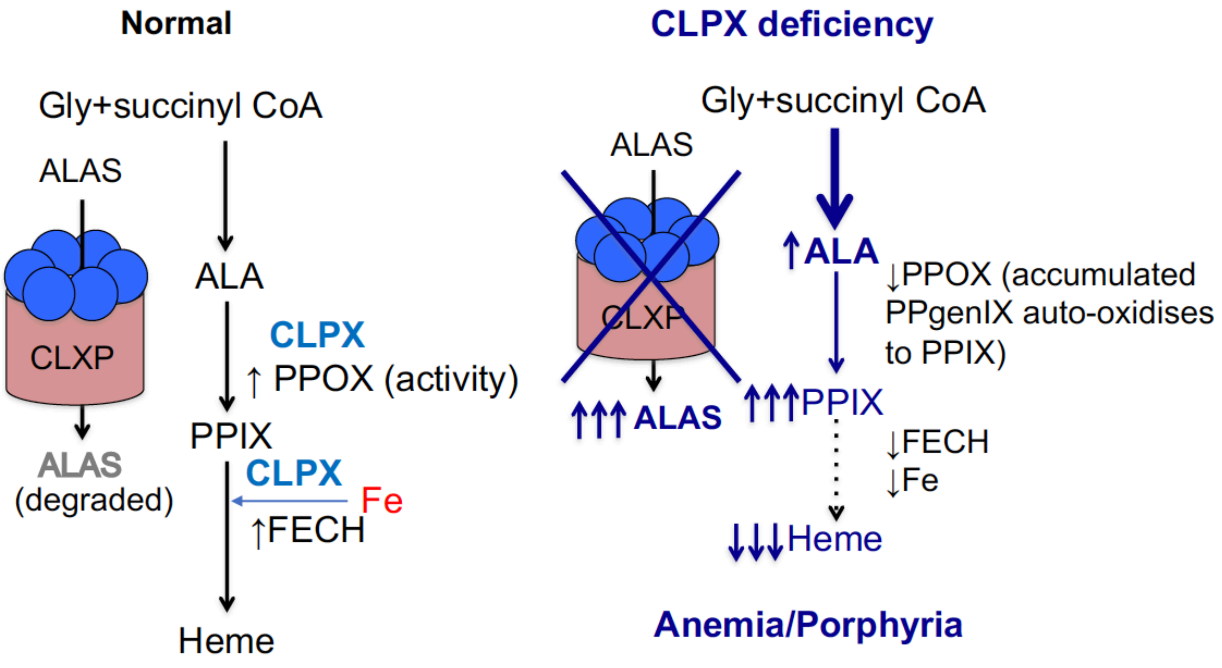


**Supplemental Figure 5.** Revised model of how CLPX regulates erythroid heme synthesis. CLPX primarily regulates ALAS2 by control of its turnover via the CLPXP protease. CLPX is required for optimal activation of PPOX, maintenance of FECH protein levels, and utilization of iron in heme synthesis (left). CLPX deficiency causes increased ALAS activity, leading to increased ALA production. CLPX deficient cells also have decreased PPOX and FECH activity, as well as an iron metabolism defect. These effects synergize to cause a large increase in PPIX levels, but a decrease in heme synthesis, potentially causing anemia and porphyria.
